# Complete genome and phylogenetic analysis of bovine papillomavirus type 15 in Southern Xinjiang dairy cow

**DOI:** 10.1101/2020.01.13.904144

**Authors:** Jianjun Hu, Wanqi Zhang, Surinder Singh Chauhan, Changqing Shi, Yumeng Song, Zhitao Qiu, Yubing Zhao, Zhehong Wang, Long Cheng, Yingyu Zhang

## Abstract

In this study, the complete genome sequence of bovine papillomavirus (BPV) type 15 (BPV Aks-02), a novel putative BPV type from a skin sample of a cow in southern Xinjiang, China was determined by collecting cutaneous neoplastic lesion, followed by DNA extraction and amplicon sequencing. The complete genome consisted of 7189 base pairs (G+C content of 42.50%) that encoded five early (E8, E7, E1, E2, E4) and two late (L1 and L2) genes. The E7 protein contained a consensus CX_2_CX_29_CX_2_C zinc-binding domain and an LxCxE motif. The nucleotide sequence of the L1 open reading frame (ORF) was related mostly (99%) to the L1 ORF of putative type BAPV-3 reference strain from GenBank. Phylogenetic analysis and sequence similarities based on the L1 ORF suggest that BPV type (BPV Aks-02) clustered with members of genus *Xipapillomavirus* as BPV15, and closely related to *Xipapillomavirus* 1.

## Introduction

Papillomaviruses (PVs) are a crescent diverse group of viruses whose genomes comprise small non-enveloped and circular double-stranded DNA virus [1–2]. The PVs have been reported worldwide to cause infection in a large variety of amniote species [2]. In cattle, bovine papilloma, also known as wart, is the most common skin tumor caused by bovine papillomavirus [3]. The BPV types and putative new PV (BAPV1 to 10) have been found by broad-spectrum detection in different places of the animals’ body, even in healthy skin [4–5]. Several studies have reported that BPV viral types have limited relationship to clinical status, type of herd, and age of animals [6–7]. This indicates that traditional clinical diagnostic techniques may not be effective methods to determine BPV types.

Recently, culture-independent molecular techniques are used to detect the PV without virus culture system [8]. Based on the sequence similarity of the highly conserved ORF L1 of papillomaviruses (PVs), the BPV types have been characterized and classified into five genera, namely *Deltapapillomavirus*, *Epsilonpapillomavirus*, *Xipapillomavirus*, *Dyoxipapillomavirus* and *Dyokappapapillomavirus* [9–10]. So far, 23 genotypes have been reported [11–12]. The BPVs 1, 2, 13, and 14 are classified in *Deltapapillomavirus* genus; BPVs 5 and 8 are grouped in *Epsilonpapillomavirus* genus; *Xipapillomavirus* genus comprised the BPVs 3, 4, 6, 9, 10, 11, 12, 15, 17, 20, and 23; BPV 7 is classed as a member of *Dyoxipapillomavirus* genus, *Dyokappapapillomavirus* genus includes BPVs 16 and 18. Other new BPV 19 and BPV 21 still have undefined genera [11–15]

In 2014, we observed a case of warty lesion from a dairy cow in Southern Xinjiang, China. Although the genomes of BPV 15 were submitted to the GenBank database, the complete genome sequence of BPV 15 has not yet been characterized. Therefore, the aim of this study was to conduct a complete genome and phylogenetic analysis of BPV type 15 in Southern Xinjiang dairy cow.

## Material and Methods

### Sample collection

In 2014, 11 warty lesion samples were collected from a dairy farm in southern Xinjiang, China. All samples were studied by L1 gene and phylogenetic analysis, but for the purpose of this publication, we focused on reporting the novel BPV 15 which was isolated from the warty lesion sample collected from one of the infected cows. The lesion was dark grey, 0.5-1 cm in diameter and located on the neck region below jaw. The sample was suspended in a 50% glycerol phosphate buffer and stored at −20 °C. The sample was named as Aks-02.

### DNA extraction

The frozen warty lesion specimens were homogenized. Total genomic DNA was extracted from warty lesions specimens using TIANamp Genomic DNA Kit (TIANGEN BIOTECH, Beijing, China) according to the manufacture’s protocol. The extracted DNAs were dissolved in 50 μl TE buffer and stored at −20 °C until used.

### PCR assay

Papillomaviral DNA was initially analyzed by PCR using the degenerate primers FAP59(forward: 5′ -TAACWGTIGGICAYCCWTATT-3′) and FAP64 (reverse: 5′ -CCWATATCWVHCATITCICCATC −3′) [16]. The expected length was approximately 480 bp.

To further characterize the complete genome sequences of the BPV Aks-02, amplification primers and sequencing primers were designed by DNAStar version 5.0 software according to available genomes in GenBank under the following accession number: AY300819, HQ612180. The primers set for amplification of the complete BAPV3 genome were BAPV-3_F (forward: 5′ -CAGTGACACCTATTCCAAGAGGTT-3′ and BAPV-3_R2 (reverse: 5′-GCATGGACCCTAAACAAGTGCAAC −3′). Details of the sequencing primers are shown in S1 Table. Both amplification and sequencing primers were synthesized by Sangon Biotech (Shanghai, China). The amplification of viral DNA by PCR was carried out base on the manufacturer’s recommendations of the LA PCR Kit (TaKaRa Biomedical Technology Beijing, China). Briefly, in a total volume of 20 μl: 2x LA Buffer 2.0 μl, dNTP (2.5 mM) 0.5 μl; Primer BAPV-3_F (10 mM) 0.4 μl; Primer BAPV-3_ R2 (10 mM) 0.4 μl; template DNA 1.0 μl; LATaq enzyme (5 U/μl) 0.4 μl; ddH_2_O 15.3 μl, which was homogeneously mixed and short spun. The expected length was approximately 7200 bp.

The PCR was performed in a thermocycler (TECHNE, TC-1500, UK) with the following cycling profile: an initial denaturation of 10 min at 94 °C, followed by 32 cycles of 30 s at 95 °C, 45 s at 56 °C and 80 s at 72 °C; and a final extension step of 10 min at 72 °C. 5 μl of PCR products were loaded on 1.5% agarose gel in Tris-acetate EDTA buffer at constant voltage (90V) for approximately 45 min, and visualized under UV light.

### Sequencing and sequence analysis

All products were purified using AxyPrep DNA Gel Extraction kits (Axygen Biotechnology, Hangzhou, China) according to the manufacturer’s instruction. The PCR amplifications of the complete genomes were sequenced bi-directionally by amplification prime and multiply sequencing primers using the Applied Biosystems 3730XL DNA Analyzer (Applied Biosystems, USA). The data of sequencing were assembled using DNAStar version 5.0 software, and submitted to Blastn search. The characteristics of the complete sequence of the PV, including predictions of putative ORF, molecular weight, motifs, and regulatory sequences were predicted using DNAStar version 5.0 software.

### Phylogenetic analysis

The complete genome sequences of L1 gene of 22 genotypes (BPV 1 to 22) from GenBank, the partial gene sequences of L1 gene of 10 putative BPV types (BAPV 1 to 10) from GenBank, were imported and aligned using Clustal W in DNAStar version 5.0 software. Phylogenetic trees were constructed from the alignment of L1 sequences using Neighbor-joining method in MEGA 6.0 software.

### Gene sequence accession number

The complete sequence of the novel PV was deposited in the GenBank database with the accession number KM983393.1. The GenBank accession numbers of these different genotype sequences used in sequence analysis and phylogenetic analysis were shown in S2 Table.

## Results and Discussion

### PCR, sequencing and sequence alignments

In this study, papillomaviral DNA from the sample of warty lesion was successfully amplified by the PCR with the degenerate primers (FAP59/FAP64). The expected length of PCR product was approximately 480 bp. The negative control with Millipore water for PCR amplification resulted in no amplified product. Based on conserved region within the L1 ORF of different genera of PVs, BPVs are divided into different genotypes. Ten putative BPV types (BAPV 1 to 10) with the partial ORF L1 nucleotide sequence were also used for sequence alignments [17–18]. Interestingly, the ORF L1 nucleotide sequence of the BPV Aks-02 was more than 99% identity with BAPV-3 (AY300819). The result revealed that BPV Aks-02 and BAPV-3 were most closely related to each other, according to difference between 2% and 10% based on L1 nucleotide sequence [1, 11]. Except for BAPV-3, present identity between the ORF L1 nucleotide sequences of the BPV Aks-02 and BPV 3, −4, −6, −9, −10 were 74.6%, 72.9%, 72.2%, 75.2% and 71.8%, respectively. Nucleotide sequence similarity of L1 gene sequence of BPV was shown in S3 Table.

Based on the results of sequence alignments analysis of L1 gene, the complete genome sequences of the identified BPVAks-02 was successfully amplified and sequenced with 17 sequencing primers, resulting in amplicons with approximately 7189 bp size (S1 Fig). The complete genomes of different BPV genotypes consisted of different ORFs and contained at least four common ORFs, there are E1, E2, L1 and L2 ORF [9, 19]. According to E1, E2, E7, L1 and L2 nucleotide sequence alignment between BPV Aks-02 and related BPV, those of BPV Aks-02 shared more than 62.2% similarity with five ORFs of *Xipapillomavirus*1 respectively. Similarly, similarities between the complete genomes of BPV Aks-02 and those of *Xipapillomavirus*1 were a maximum of 67.5-72.3% identity within five different genera (Table 1). Therefore, our results suggested that the BPV Aks-02 should be classified in *Xipapillomavirus*1.

**Table 1.**
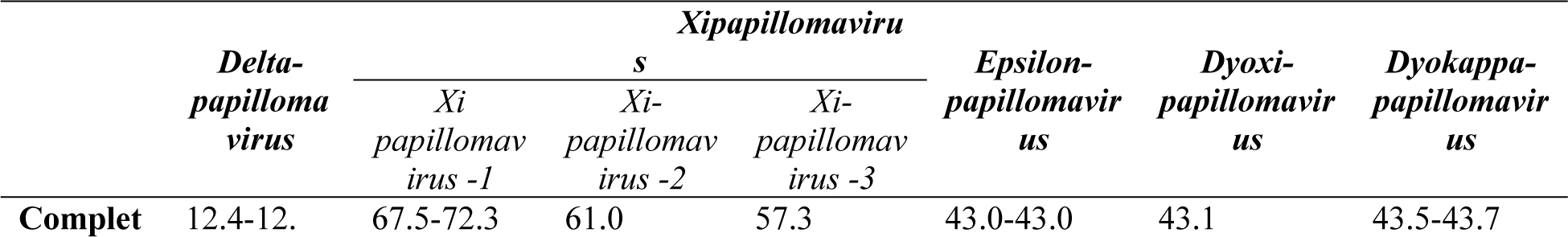
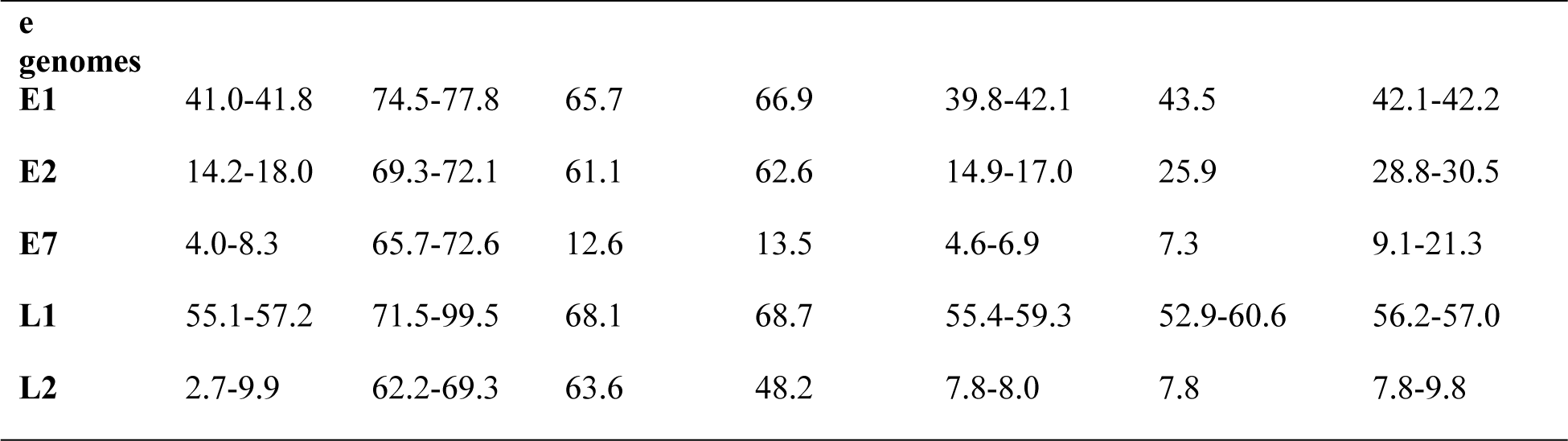
Nucleotide sequence similarity (%) between BPV Aks-02 and related BPV.

### Phylogenetic analysis

The phylogenetic tree of L1 ORF sequence was constructed with 32 BPVs using Neighbor-joining method in MEGA 6.0 software. According to the International Committee on Taxonomy of Viruses (ICTV) new *Papillomavirus* taxonomy proposal, the *Xipapillomavirus* genus includes three papillomavirus species: *Xipapillomavirus*1 (BPV3, 4, 6, 9, 10, 11, 15 and 23)*, Xipapillomavirus*2 (BPV12), *Xipapillomavirus*3 (RtPV2), respectively [11, 14, 15]. As evident from the phylogenetic tree, the BPV Aks-02 was more closely related to BAPV3-AY300819 of the species *Xipapillomavirus*1, classified in the *Xipapillomavirus* genus (Fig 1). Therefore, the phylogenetic analysis indicated that the BPV Aks-02 appeared in the same cluster as BAPV3 within the genus *Xipapillomavirus*1. Additionally, the new putative BPV types BAPV 1 to 10 were classified in different genera in the family *Papillomaviridae*. However, other two new BPV types, namely, BPV 19 and BPV 21 are still unclassified genus, which requires further research [11, 14, 15].

**Fig 1.**
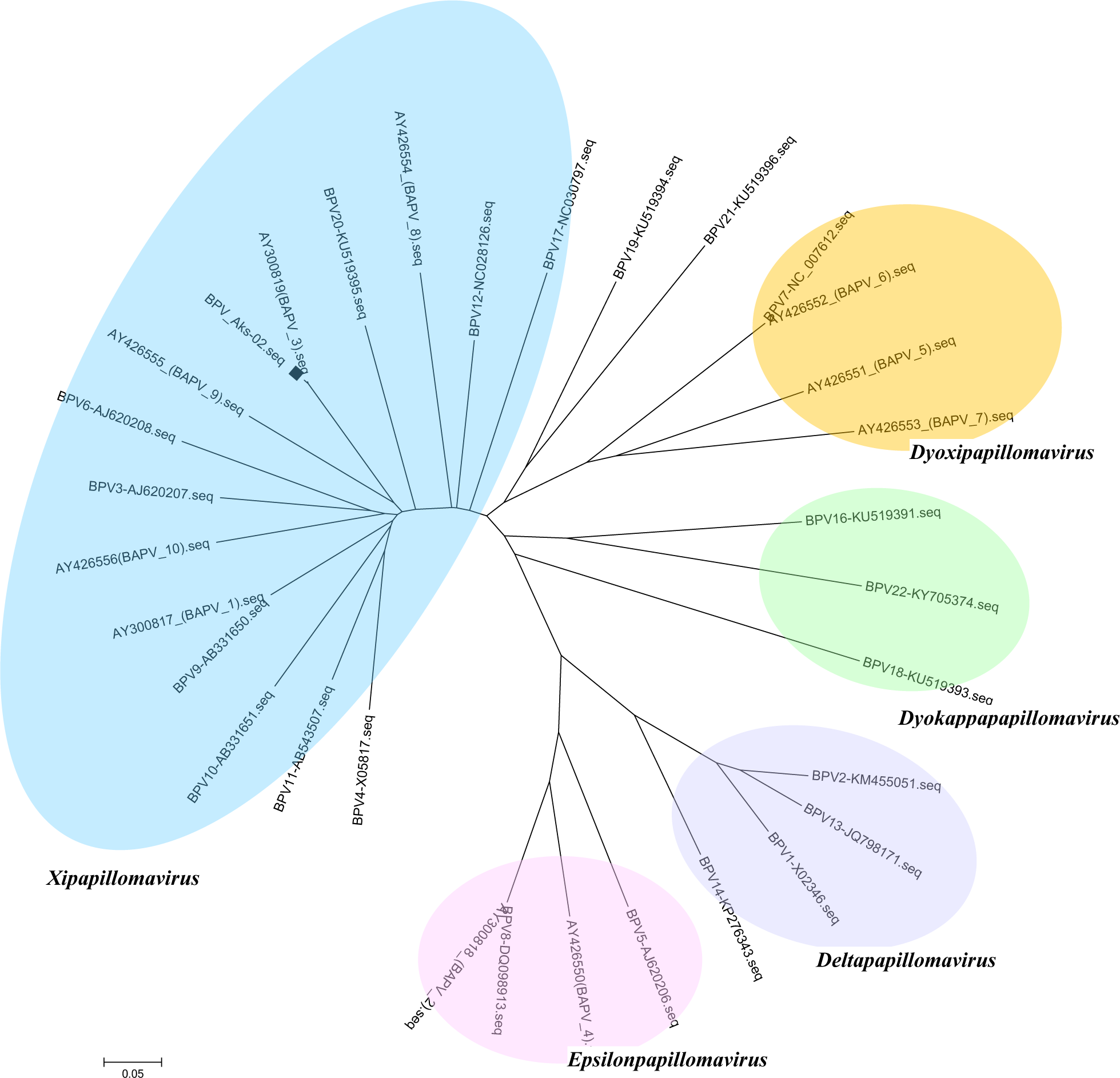
Phylogenetic tree based on the papillomavirus L1 gene. The phylogenetic tree was inferred using the neighbor-joining method based on an L1 nucleotide sequence alignment of 31 BPVs and BPV Aks-02 (the BPV Aks-02 is indicted with a black diamond). The tree was divided into the previously determined genera *Xipapillomavirus, Epsilonpapillomavirus*, *Deltapapillomavirus*, *Dyokappapapillomavirus*, *Dyoxipapillomavirus*, and unassigned PV genus (BPV19, BPV21)

### The complete BPV 15 genome structure

The complete genome structure of the BPV Aks-02 was predicted to contain 7189 bp with a GC content of 42.5% and consist of five early (E8, E7, E1, E2, E4) and two late (L1 and L2) ORFs. The long control region (LCR) located between the early and late ORFs of the circular virus. The E4 ORF was embedded within E2. E7 and E1, E1 and E2, were partial overlapping. The BPV Aks-02 was lack of characteristic E6 ORF [1]. The predicted ORFs and characteristics of genomes structure are shown in Fig 2 and Table 2.

**Table 2.** Main features of the proposed BPV genome.

|  | Position | Length (nt) | Length (aa) | Molecular mass (KDa) | Isoelectric Point |
| --- | --- | --- | --- | --- | --- |
| **LCR** | 1-473 | 473 | - | - | - |
| **E8** | 474-602 | 129 | 42 | 4858.97 Daltons | 5.411 |
| **E7** | 808-1,110 | 303 | 100 | 11140.66 | 3.938 |
| **E1** | 1,100-2,929 | 1830 | 609 | 69496.13 | 6.556 |
| **E2** | 2,871-4,085 | 1215 | 404 | 45699.32 | 9.723 |
| **E4** | 3,391-3,819 | 429 | 142 | 16301.39 | 5.938 |
| **L2** | 4,099-5,673 | 1506 | 501 | 57205.94 | 7.786 |
| **L1** | 5,684-7,189 | 1575 | 524 | 57317.08 | 4.759 |

**Fig 2.**
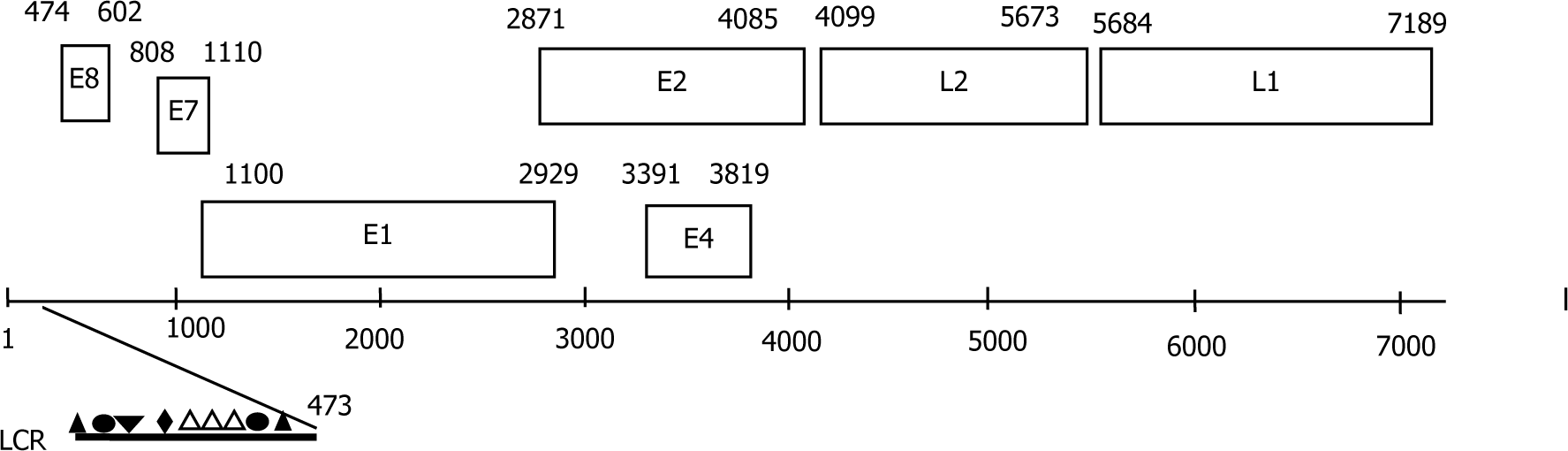
Complete genome structure analysis of the BPV Aks-02. The boxes represent the early (E) and the late (L) ORFs. The long control regions (LCR) encompassing nucleotides 1-473 ▲, E1 binding sites (E1BS); ●, Polyadenylation signals; ▼, Nuclear factor binding sites (NF-1); ◆, TATA box of the viral promoters; △, E2 binding sites (E2BS).

### The long control region

LCR known as the non-coding region (NCR) or up-stream regulatory region (URR), was located at nt 1 to 473 between the stop-codon of L1 and the start-codon of E8 in the BPV Aks-02. Many sites were found in the early region of complete genome. Most PVs possess some of the transcription factor binding sites (TFBS) in the LCR, such as E2BS, E1BS, NF-1 [20–23]. LCR of the BPV Aks-02 contained one E1BS (AACAAT at nt 28-31), two canonical E2BS (ACCGN4CGGT at nt 322-333, nt 410-421), one non-canonical E2BS (ACC-N6-GGT at nt 395-406), respectively. E2BS are essential for activating or repressing the transcription of viral genome [24]. One modified E1BS (TAACAA at nt 366-371) was presented in the LCR. It also contained only one polyadenylation (polyA) sites (AATAAA at nt 81-86) and two TATA boxes (TATAAAA at nt 261-267, nt 425-431) in the LCR. Nuclear factor binding sites NF-1 (TTGGCA at nt 113-118) were also located in the LCR.

### BPV 15 early region

The early region of the BPV Aks-02 includes 5 ORFs, such as E8 (128 bp), E7 (302 bp), E1 (1829 bp), E2 (1214 bp) and E4 (428 bp). The early region mainly encodes the non-structural viral proteins for initiation of virus replication [2, 14]. The BPV Aks-02 genome included a putative E8 gene in the early region [1] [24]. An important element of E8 protein was presented to be the conserved C-terminal amino acids L_X_GWD and repeated sequences TGTCAACTGT (nt 568-577). The BPV Aks-02 E8 ORF shared 65.1% identity of nucleotide sequence with BPV3 E8 ORF (AF486184.1) [11].

The pRbBD found in the E7 protein of most PVs members play an important role in interfering with host cell cycle and the immortalization and transformation of the host cell [11, 14, 25]. The BPV Aks-02 E7 ORF encoded a 100 aa protein and contained a conserved retinoblastoma tumor-suppressor protein-binding domain (pRbBD: L_X_C_X_E). Furthermore, one consensus zinc-binding domain (ZnBD: C_X2_C_X29_C_X2_C) was located in the C-terminal region. The CXXC motifs were separated by 29 aa residues.

The E1 and E2 of the early region in all PV are necessary for genome transcription and replication [14]. The E1 ORF coded about 609 aa as the largest protein of BVP 15. The conserved ATP-binding site of ATP-dependent helicase (G_X4_GKS) is located in C-terminal region of E1. There were two modified E1BS (TAACAA) at 1357-1362, nt 1540-1545 in the E1 ORF. Additionally, the E1 ORF contains a cyclin interaction RXL motif (the native KRRLL recruitment motif), which is essential for the initiation of papillomavirus replication and could be a potential target for developing therapeutic agents or vaccine development [14, 20]. There was a lack of a leucine-zipper domain (LX_6_LX_6_LX_6_LX_6_L) in the E2 protein of the BPV Aks-02 [26]. The Putative E4 ORF showed a start codon and nested within the E2 ORF, which is 71.8% similar to BPV 3 (AF486184.1).

### BPV 15 late region

The BPV viral capsid is mainly consisted of the major L1 and minor L2 late proteins [11]. The late region of the BPV Aks-02 was predicated to contain major capsid protein L1 and minor capsid protein L2. There were two E1BS (AACAAT at nt 5439-5444, nt 6917-6922) and one modified E1BS (TAACAA at nt 5438-5443) in the late region. For the late viral mRNA transcription, polyadenylation signals PolyA (AATAAA) were identified at nt 7040-7045 of the L1. In addition, Polyadenylation signals PolyA (ATTAAA instead of AATAAA) was founded at nt 4093-4098 between the stop-codon of the E2 and the start-codon of L2 ORF. There were one canonical E2BS (ACC-N6-GGT at nt 7126-7137) in the L1 gene and one non-canonical E2BS (ACC-N7-GGT at nt4308-4320) in the L2 gene. Nuclear factor binding sites (NF-1: TTGGCA at nt 5278-5283) were also found in the L2 ORF. The L1 protein was presented a high proportion of positively charged residues (K or R) in the C-terminal end.

## Conclusion

In this study, we detected the BPV genotype from a neck wart of dairy cow and present the complete genome sequences and genome characterization of BPV 15. Based on these genetic features, it is suggested that BPV 15 classified viral type in the *Xipapillomavirus* genus, well aligns in *Xipapillomavirus* genus 1 branch. These data may contribute to the understanding of the classification, genomic evolution and epidemiology of BPV.

## Supporting information

**S1 Table.**
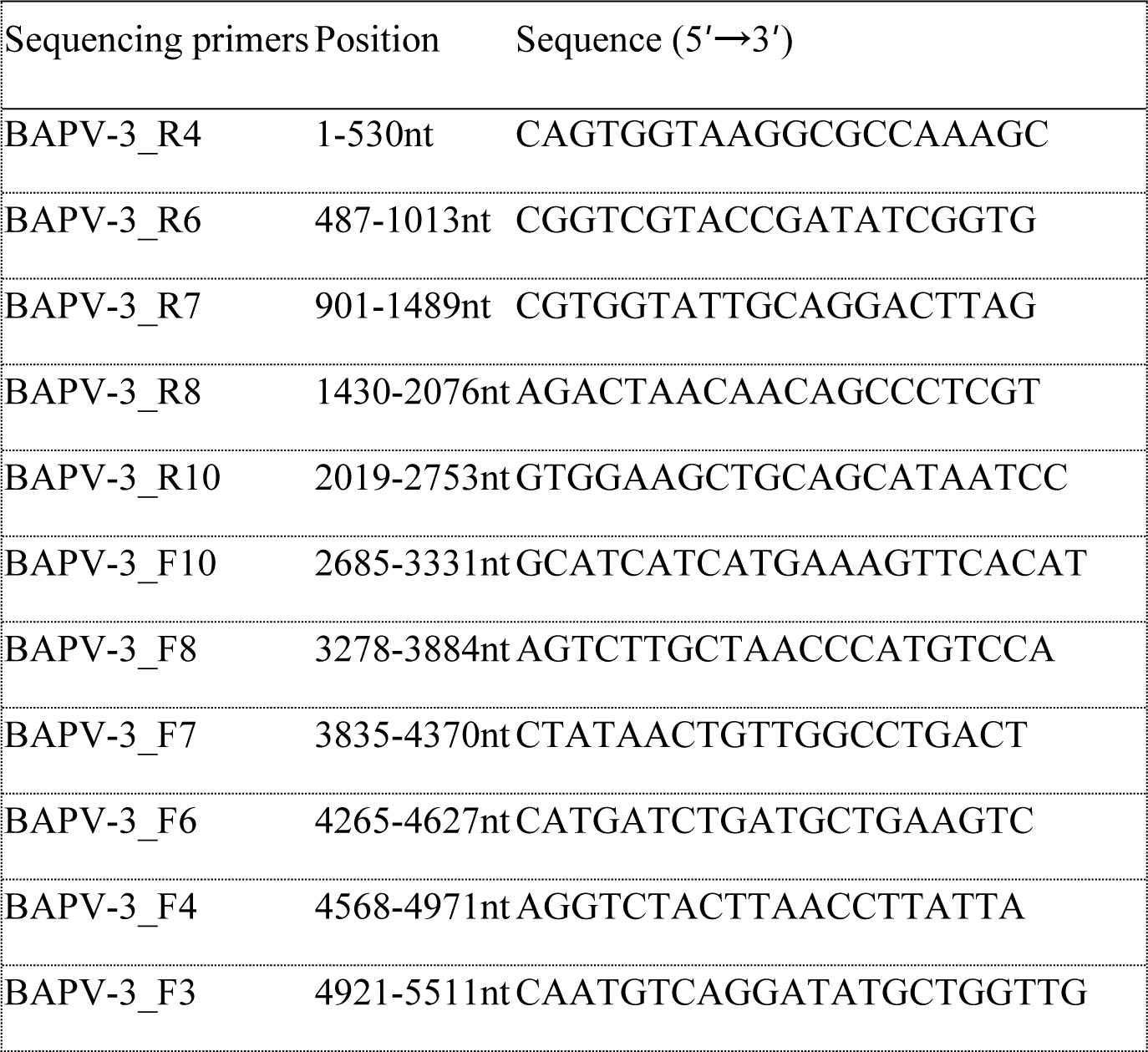
Sequencing primers.

**S2 Table.**
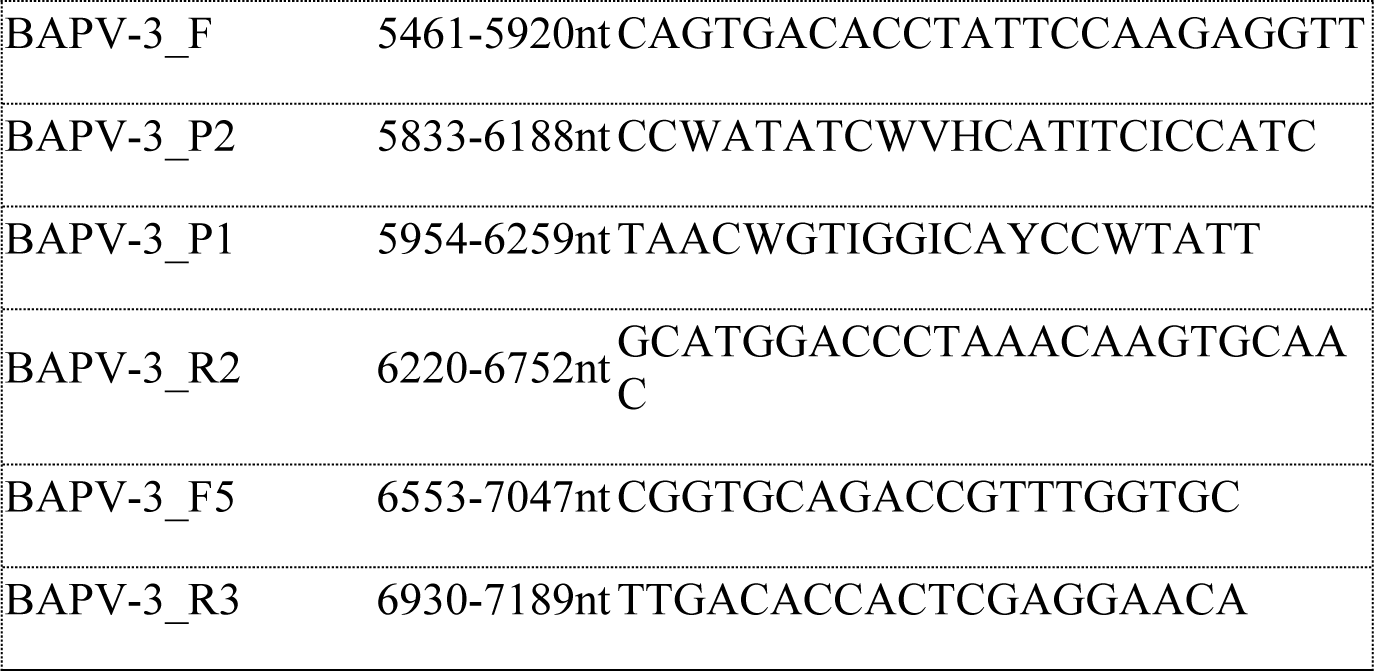
Reference strains. All reference strains in this study were downloaded from GenBank (http://www.ncbi.nlm.nih.gov/), as shown in Table with the respective GenBank accession numbers.

**S3 Table.** Homology analysis of L gene sequence of BPV

| Scientific<br>Abbreviation | genus | accession number |
| --- | --- | --- |
| BPV1 | *Deltapapillomavirus* | X02346 |
| BPV2 | *Deltapapillomavirus* | KM455051.1 |
| BPV3 | *Xipapillomavirus* | AF486184.1 |
| BPV4 | *Xipapillomavirus* | X05817.1 |
| BPV5 | *Epsilonpapillomavirus* | AF457465 |
| BPV6 | *Xipapillomavirus* | AJ620208.1 |
| BPV7 | *Dyoxipapillomavirus* | NC_007612 |
| BPV8 | *Epsilonpapillomavirus* | DQ098913 |
| BPV9 | *Xipapillomavirus* | AB331650 |
| BPV10 | *Xipapillomavirus* | AB331651 |
| BPV11 | *Xipapillomavirus* | AB543507 |
| BPV12 | *Xipapillomavirus* | NC_028126.1 |
| BPV13 | *Deltapapillomavirus* | JQ798171 |
| BPV14 | *Deltapapillomavirus* | KP276343 |
| BPV15 | *Xipapillomavirus* | KM983393.1 |
| BPV16 | *Dyokappapapillomavirus* | KU519391 |
| BPV17 | *Xipapillomavirus* | NC_030797.1 |
| BPV18 | *Dyokappapapillomavirus* | KU519393.1 |
| BPV19 | undefined | KU519394.1 |
| BPV20 | *Xipapillomavirus* | KU519395.1 |
| BPV21 | undefined | KU519396.1 |
| BPV22 | *Dyokappapapillomavirus* | KY705374 |
| BAPV 1 | *Xipapillomavirus* | AY300817 |
| BAPV 2 | *Epsilonpapillomavirus* | AY300818 |
| BAPV 3 | *Xipapillomavirus* | AY300819, HQ612180 |
| BAPV 4 | *Epsilonpapillomavirus* | AY426550 |
| BAPV 5 | *Dyokappapapillomavirus* | AY426551 |
| BAPV 6 | *Dyokappapapillomavirus* | AY426552 |
| BAPV 7 | *Dyokappapapillomavirus* | AY426553 |
| BAPV 8 | *Xipapillomavirus* | AY426554 |
| BAPV 9 | *Xipapillomavirus* | AY426555 |
| BAPV 10 | *Xipapillomavirus* | AY426556 |

**S1Fig.**
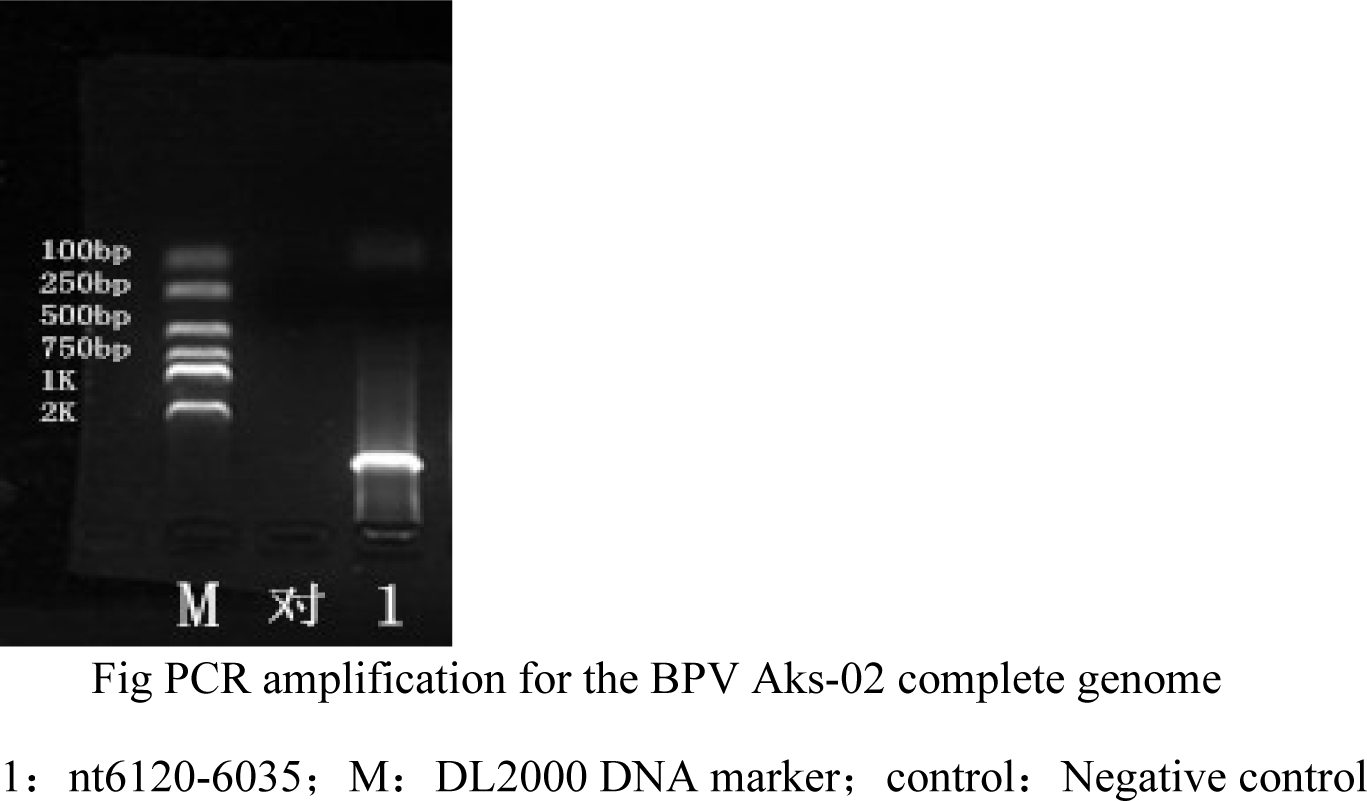
PCR amplification fragment. A PCR amplification fragment was obtained and the results of the 1.5% agarose gel electrophoresis showed that the fragment size was 7189 bp.

## Acknowledgments

We thank National Nature Science Foundation of China for financial support.

